# An Extensive Sequence Dataset of Gold-Standard Samples for Benchmarking and Development

**DOI:** 10.1101/2020.12.11.422022

**Authors:** Gunjan Baid, Maria Nattestad, Alexey Kolesnikov, Sidharth Goel, Howard Yang, Pi-Chuan Chang, Andrew Carroll

## Abstract

Accurate standards and extensive development datasets are the foundation of technical progress. To facilitate benchmarking and development, we sequence 9 samples, covering the Genome in a Bottle truth sets on multiple instruments (NovaSeq, HiSeqX, HiSeq4000, PacBio Sequel II System) and sample preparations (PCR-Free, PCR-Positive) for both whole genome and multiple exome kits. We benchmark pipelines, quantifying strengths and limitations for sequencing and analysis methods. We identify variability within and between instruments, preparation methods, and analytical pipelines, across various sequencing depths. We discuss the relevance of this variability to downstream analyses, and strategies to reduce variability.

## Introduction

Accurate standards and extensive development datasets are the foundation of technical progress. In genomics, the Platinum Genomes and Genome in a Bottle have produced truth sets for small variants [1–3] and structural variants [4], and methods for their comparison [5, 6] and curation [7]. These are critical to development and assessment of read mapping [8, 9], variant calling [10–13], phasing [14], and genome assembly [15–18].

Genome in a Bottle uses multiple sequencing technologies at high coverage and accuracy in order to create the most comprehensive and correct truth set for NA12878/HG001 from 1000 Genomes [19], and two trios (HG002-HG003-HG004 and HG005-HG006-HG007) from participants in the Personal Genomes Project [20]. While the Genome in a Bottle datasets are large and high quality, most are from 2016 and use sequencing instruments (HiSeq2500) and protocols that are not fully representative of common use today.

Here, we release a sequence dataset generated to broadly represent common whole genome sequencing (WGS) preparations (PCR-Free and PCR-Positive), multiple types of whole exome sequencing (WES) capture kits (Agilent v7, IDT, Nextera, and Truseq) and instruments (Illumina NovaSeq and HiSeqX), for nine samples: the HG002-HG003-HG004, HG005-HG006-HG007, and NA12878/HG001-NA12891-NA12892 trios.

To facilitate development and comparisons at the range of typical use, we have pre-mapped the sequencing runs with BWA MEM [21] both GRCh37 and GRCh38 [22]. To allow comparisons along a range of typical coverages, we release samples at 40x, 30x, and 20x mapped coverage for WGS and 100x, 75x, and 50x on-target coverage for the exomes.

We characterize variability in sequencing runs within and between sequencing platforms, coverages, sample preparation and exome capture kit, and common variant calling pipelines.

To contribute to efforts to expand the truth set into difficult regions of the genome, we also ordered PacBio HiFi [23] sequencing of HG003, HG004, HG006, and HG007. We characterize coverage and variant calling aspects of these runs, contrasting them with Illumina WGS.

## Results

We contracted HiSeqX and NovaSeq WGS sequencing from Novogene for both PCR-Free and PCR-Positive preparations at 50x coverage. We contracted exomes on HiSeq4000 and NovaSeq with Agilent v7, IDT-xGen kits. In total, there are 36 WGS and 54 WES samples. All Illumina sequencing is paired-end with read lengths of 151 bp.

These samples were mapped using BWA MEM[21] and were deduplicated with Picard[24]. We assessed the proportion of duplicate, unmapped, and uniquely mapped reads with samtools flagstat[25], and coverage across the genome and capture regions using mosdepth[26].

### Coverage, Mapping, and Target Capture Variability

For WGS samples, we did not observe a large difference in raw coverage or mapped coverage between NovaSeq and HiSeqX, or PCR-Free and PCR+ (Figure 1 top). The proportion of duplicate reads was 13.7% ± 3.7%, with one outlier sample at 26.3%. The proportion of unmapped reads was 0.7% ± 0.8%.

**Figure 1.**
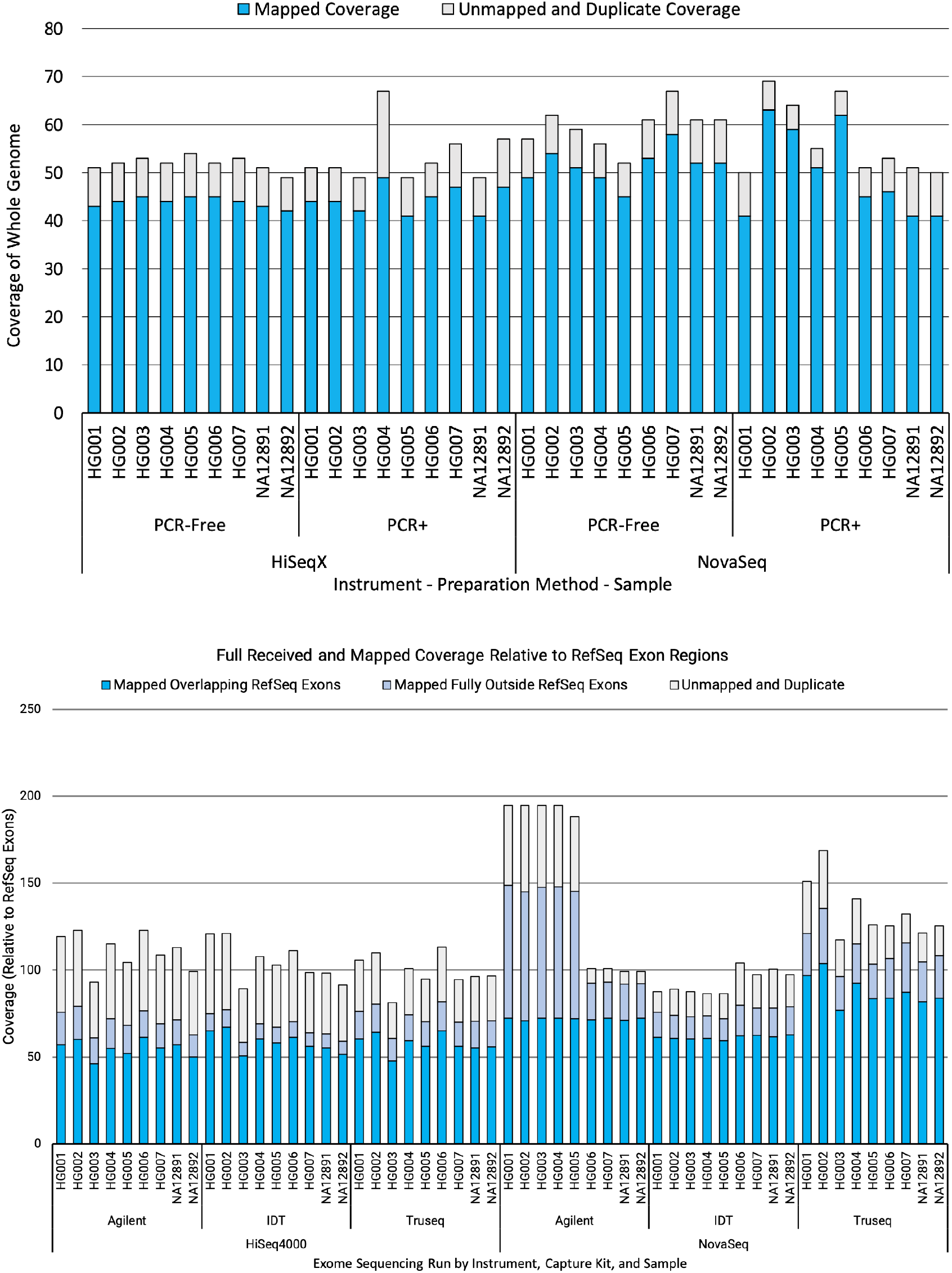
Coverage of WGS and WES samples in this study. Raw coverage received and the coverage of deduplicated, mapped reads for WGS samples on GRCh38 with ALT contigs (top). For WES, whether coverage overlaps RefSeq is also shown (bottom).

For WES, received coverage was more variable between kits and runs. However, the coverage overlapping GRCh38 RefSeq exons was more consistent, with a range of 59x-87x coverage (71x ± 8x) (Figure 1 bottom). A substantial amount of coverage was mapped outside of exons.

To facilitate community use on standardized, processing-ready samples, we generated serial downsamples of the WGS at 50x, 40x, 30x, and 20x uniquely mapped coverage, and of the exomes at 100x, 75x, and 50x coverage of the kit capture regions. These are mapped to GRCh37 and GRCh38 with ALT contigs in an ALT-aware manner. At higher coverages, some samples are not present due to insufficient coverage. In total, this covers 246 WGS BAM files and 218 WES BAM files. These files are available for public download (see data availability) without access control or restriction. Download URLs for FASTQs (Supplementary File 1), BAMs (Supplementary File 2), and VCFs (Supplementary File 3) are included.

To better understand the uniformity of sequencing across multiple benchmark samples, we investigated the per-base coverage across each of the samples at 30x median coverage. The WGS samples had roughly 50% of the genome coverage at 30x, as expected. In terms of mapped coverage, there were not notable differences between PCR-Free and PCR+ preparations. However, 4 PCR-Free NovaSeq and 2 PCR-Positive NovaSeq samples had a noticeable difference in coverage profile (Figure 2A). We stratified by various GA4GH regions [27], revealing a large difference in regions with GC content <25% (covering ~140 MB or 5% of the genome) (Figure 2 top right). The difference was reduced, but still visible in regions overlapping RefSeq [28] exons. There was more per-run variability in regions with GC content >75%, with the 6 previously identified NovaSeq samples having slightly higher fraction of the genome covered (Supplementary Figure 1 top right).

**Figure 2.**
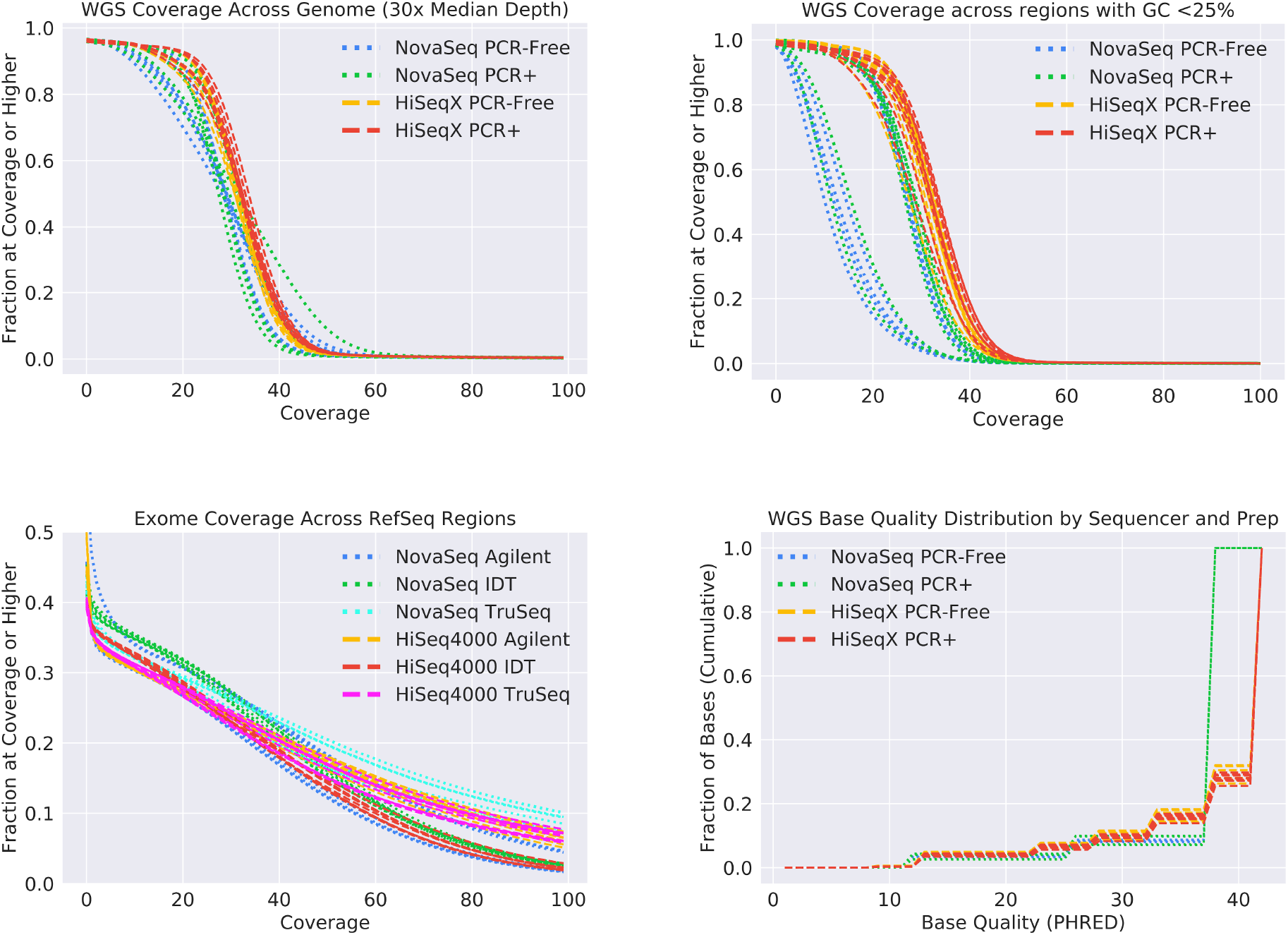
Genome coverage and read quality attributes for samples in this study. The fraction of the genome in each WGS sequencing run covered to a least a given coverage (top left). Restricting this analysis to GA4GH regions with GC content less than 25% reveals greater variability by instrument for WGS (top right). The fraction of the genome overlapping RefSeq exon regions for exomes covered to at least a given coverage (bottom left). Exome samples used are at 50x median on-target coverage. The base quality distribution by sequencer and preparation for WGS (bottom right).

We also observed that in tandem repeat and homopolymer regions, NovaSeq samples have a consistent, but small reduction in the fraction of the genome covered compared to HiSeqX samples (Supplementary Figure 1 bottom left). For exomes, we investigated coverage of RefSeq [28] exons. We found greater variability between samples due to large differences in capture efficiency (Figure 1 bottom), though to the extent that a trend is discernible, the type of kit used had a more noticeable effect on coverage of the RefSeq exons.

We investigated the distribution of base-qualities between these sequencing runs. The NovaSeq has 4 bins of quality values, while HiSeqX and HiSeq4000 use 8 bins of values. For WGS, samples had a high degree of consistency in their base quality distribution profile. Despite the different binning schemes, the distribution of base qualities is similar for NovaSeq and HiSeqX/HiSeq4000 (Figure 2 bottom right). The exome sequences were similar, though there was a higher degree of variability and lower average base quality in the exomes sequenced from HiSeq4000 runs on the Agilent kit (Supplementary Figure bottom right).

### Accuracy of Variant Calling Pipelines for WGS

In order to understand the impact of choices for instrument, preparation, and coverage, we performed variant calling with several different widely-used, open-source methods: GATK4 [11], Strelka2 [29], Octopus[30], and DeepVariant v1.0 [12]. We assessed the accuracy of the pipelines against the Genome in a Bottle truth sets for HG001-HG007[1]. For HG002, HG003, and HG004, we use the v4.2, which is more comprehensive, accurate, and built natively on GRCh38 [27]. For HG001, HG005, HG006, and HG007, we compare using the v3.3.2 truth set on GRCh37, since the truth set was generated on GRCh37 and lifted over to GRCh38.

To visualize the variability in performance, we plotted the precision and recall for both NovaSeq and HiSeqX on each of HG002-HG004 for PCR-Free (Figure 3 top) and PCR-Positive (Supplementary Figure 2) samples at 20x, 30x, and 40 coverage. For SNP accuracy (Figure 3 top left), accuracy clearly separates by analysis method, with DeepVariant having high precision and recall, Strelka2 having high precision, and Octopus having having higher precision than GATK, with recall between Strelka2 and DeepVariant. For Indels (Figure 3 top right), the choice of method has a larger impact than coverage. For Indels, the effect of the analytical method remains pronounced, but coverage has a more pronounced impact on the results.

**Figure 3.**
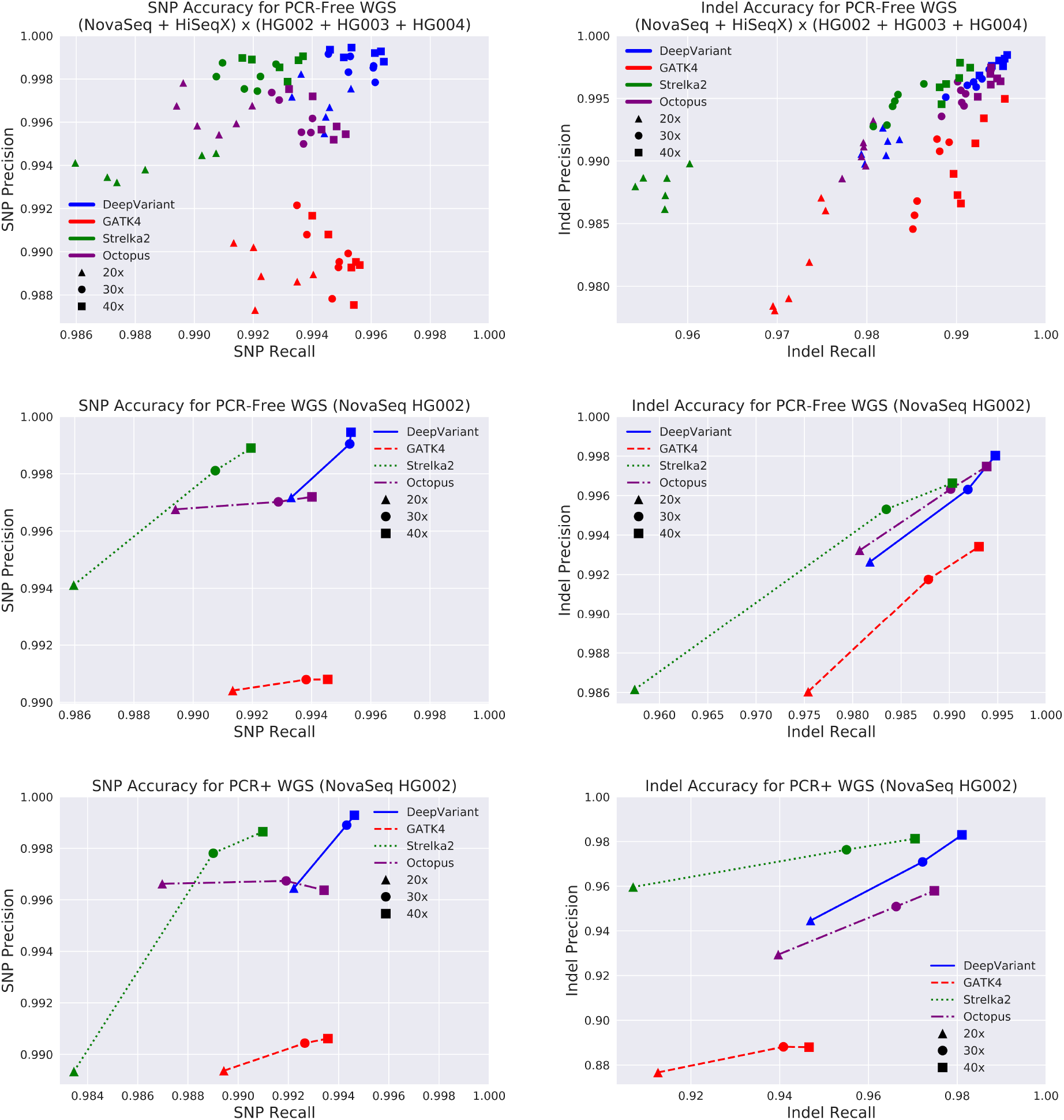
Variant Calling Accuracy for WGS Samples. Precision and Recall for SNP (left) and Indel (right) for WGS sequence runs, compared with the v4.2 GIAB truth set on BAM files mapped to GRCh38. All accuracies are for chromosome20, which is not used for training DeepVariant. Accuracies across multiple PCR-Free runs of HiSeqX and NovaSeq for each of HG002, HG003, and HG004 at 20x, 30x, and 40x coverage (6 points at each coverage, 18 total runs). Four different callers were run on the same samples, coded by color: DeepVariant, GATK4, Strelka2, Octopus (top). Accuracies for a single sample (NovaSeq HG002) for PCR-Free WGS (middle) and for PCR+ (botton). Note the higher dependence of coverage for Indel calling, and the different axis scale for Indel PCR-Free and PCR+, indicating the greater impact of PCR on Indel calling accuracy.

Preparations which include PCR achieved similar SNP accuracies to PCR-Free, but have much lower Indel accuracies (Supplementary Figure 2 bottom, Figure 3 bottom right), likely reflecting increased error around homopolymer sites. Some of the analytical methods are more robust to these increased errors, likely due to specific effort to model this error. These methods are Strelka2, which specifically includes a PCR Indel error model, and DeepVariant, which includes PCR-Positive data in its training set. The greater effect of coverage observed for Indel accuracy, sequencing to higher depth allows coverage to partially mitigate inclusion of PCR.

### Accuracy of Variant Calling Pipelines for Exome

The exome sequencing spans three different capture kits, which each target different regions. We performed variant calling with each of the pipelines (using exome-specific tool settings where appropriate, for example using the --exome flag for Strelka2 and the exome model for DeepVariant). For GRCh37, we performed calling on the union of all capture regions and RefSeq genes. For GRCh38, we performed calling on the RefSeq exons regions. We padded each region with 100bp of each included region, to maximize calls available to others. However, the comparisons here use only directly included regions. We plot NovaSeq and HiSeq4000 for HG002-HG004 using the intermediate coverage point (75x) for the RefSeq exon regions (Figure4 top) and the shared intersection of capture regions (Figure 4 bottom).

**Figure 4.**
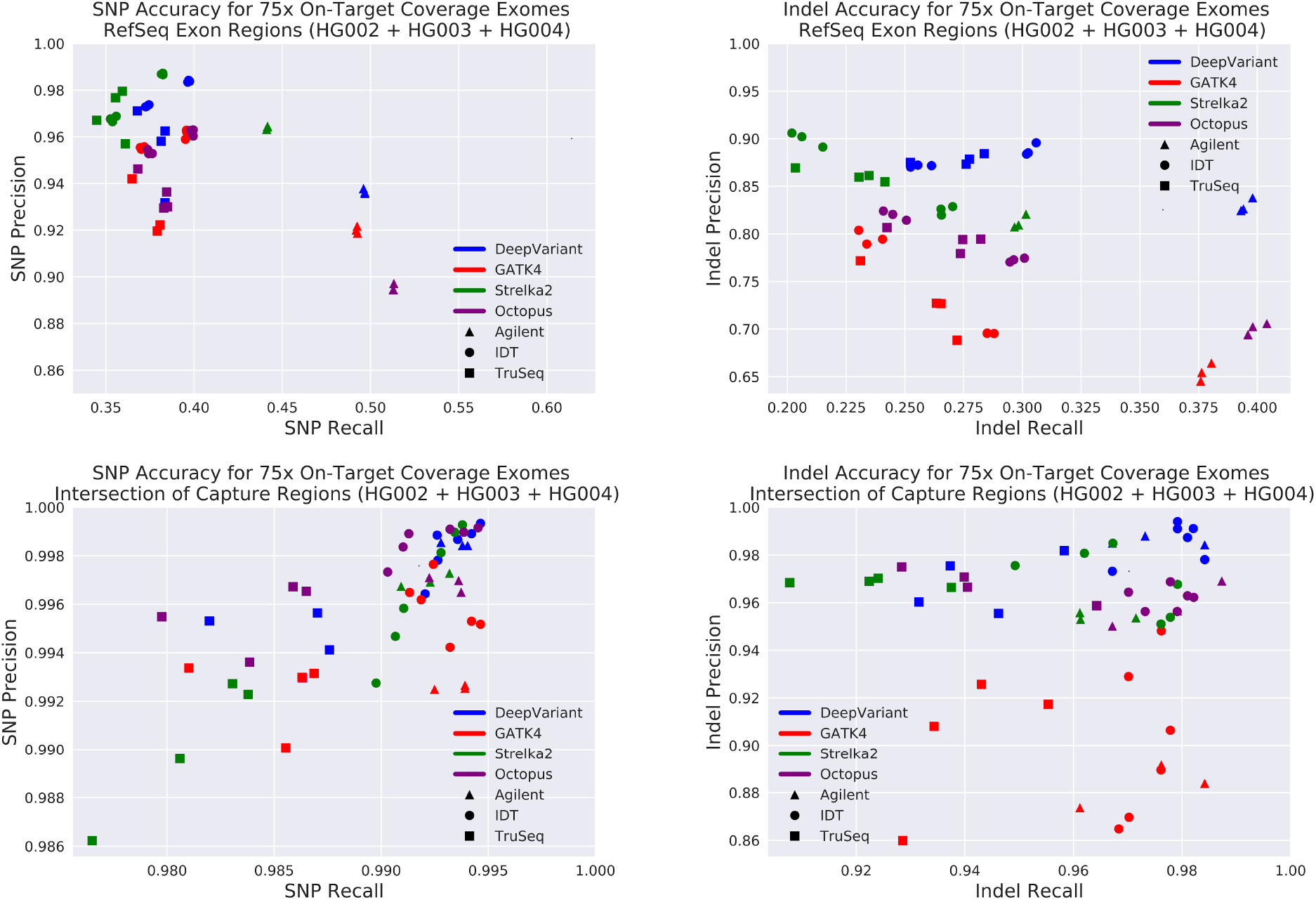
Variant Calling Accuracy for Exome Samples in RefSeq and Capture Regions. Precision and Recall for SNP (left) and Indel (right) for exome sequence runs, compared with the v4 GIAB truth set. All accuracies are across the full genome in order to have enough variants for a stable comparison. Four different callers were run on the same samples, coded by color: DeepVariant, GATK4, Strelka2, Octopus, with capture kits coded by shape for: Agilent, IDT, TruSeq. The comparison uses a restricted set of genome regions: all RefSeq exon regions in GRCh38, covering ~136 MB (top), and the intersection of capture regions for the three exome kits, covering ~32 MB (bottom). In both cases, all kits are compared on the same regions.

Compared to the WGS results, accuracy on exomes is lower, consistent with other studies that used Sanger validation[31]. For both all RefSeq exons and the share capture regions, Indel accuracy was much lower. This may reflect the interaction between the greater coverage dependence for Indel accuracy and variability in capture efficiency, and/or the preparation of the exome samples.

Recall across all RefSeq exons is much lower than for shared capture regions, likely reflecting lower or absent coverage outside capture regions. Precision is lower as well, for both SNPs and Indels. This may be due to coverage variability from more difficult to capture regions that would be present in one kit, but not the others. There is a clear separation of accuracy by analytical method for Indel calling, especially across the RefSeq exons, with DeepVariant having higher accuracy. This could be explained by DeepVariant having better accuracy at lower coverages, and/or DeepVariant learning some of the error profiles specific to exome sequencing. (DeepVariant was not trained with any of these exome samples or these kits).

In comparing the performance of the different kits to each other, the best-performing kit depends on the type of results desired. Across all capture regions, the Agilent kit allows recall or a larger number of variants, consistent with another study[32]. However, the on-target comparison indicates that the IDT kit has a higher accuracy on sites shared in each of the capture regions. Strategies such as those employed for the exome sequencing of UK Biobank, which supplement the IDT exome with additional capture probes could be one strategy for achieving a good balance of on-target accuracy and breadth of capture[33].

### Contributions to Error across All Factors

Across all combinations to sequence and analyze a WGS run for all samples with a truth set, there are 336 different permutations, and 504 different exome permutations. This raises a tension between presenting the breadth and variability across all combinations, and presenting a small number of comparisons which are easier to visualize and compare. The analysis in prior sections presented a subset of the data, with both grouped and individual visualizations.

We sought a metric that could compare the factors contributing to analysis quality across all of the permutations. To do this, we binned the factors into instrument, preparation, variant caller, and coverage. We calculate the mean and standard deviation of SNP and Indel errors, and calculate the number of standard deviations from the mean each sample with a given property is. We use v3.3.2 Truth set and GRCh37 for HG001, HG005-HG007, and the v4.2 Truth set and GRCh38 for HG002-HG004., performing the standard deviation calculations separately for v4.2 and v3.3.2 samples, due to the differences in number of variants between the truth sets. This allows a common metric for accuracy (standard deviations from mean) across all samples

Comparisons for WGS (Figure5 top), exomes evaluated across RefSeq exons (Figure5 bottom) and the shared capture regions (Supplementary Figure 4) indicate which components contribute to pipeline accuracy and consistency. As expected, increasing coverage increases the accuracy of all pipelines, with the largest increase coming from increasing from the lowest coverage points. Increasing coverage also increased the consistency between runs. PCR-Positive WGS preparations have lower accuracy than PCR-Free, with a more pronounced effect on Indels than SNPs. For WGS pipelines, NovaSeq runs had a lower SNP accuracy as opposed to HiSeqX runs, while NovaSeq runs had a higher Indel accuracy than HiSeqX runs. The magnitude of accuracy change for coverage from the lowest coverage point to the highest coverage point was similar to the magnitude of change from the most least accurate caller to the most accurate caller, and for Indels the difference from PCR-Positive to PCR-Free.

**Figure 5.**
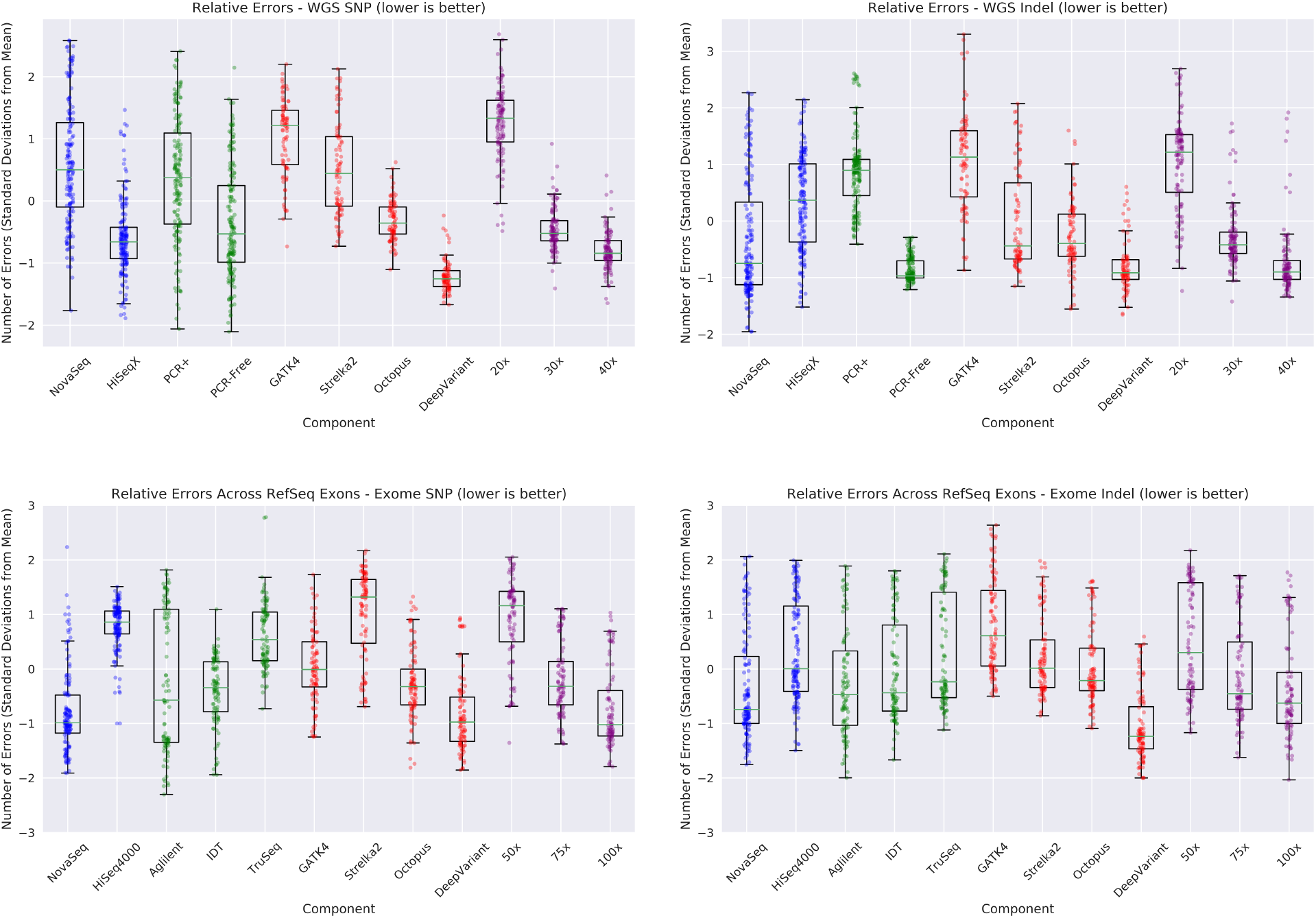
Contributions of Components to Variant Calling Pipeline Accuracy. Measuring the contributions of component: instrument (red), preparation/kit (green), variant caller (red), and coverage (purple) to SNP (left) and Indel (right) errors in variant calling. The mean and standard deviation for error is calculated, and for each sample the number of standard deviations from the mean of errors is plotted. For example, every dot in the NovaSeq column is a run using NovaSeq data where there is another run with all of the same properties except the use of HiSeqX in the HiSeqX column. When a dot is below y-axis=0, it means the use of that factor reduces error relative to the “average” run. This is shown for WGS pipelines (top) and for exome pipelines (bottom).

For exomes, the consistency of results across all runs was lower than for WGS. For both SNPs and Indels, the NovaSeq instrument runs had higher accuracy than the HiSeq4000 ones. The difference in accuracy from the lowest coverage to the highest coverage point was again similar to the difference between the least accurate and most accurate variant calling method. For exome kits, Aglient had a slightly higher accuracy when evaluated across all of RefSeq (Figure 5 bottom), while IDT had a slightly higher accuracy on the shared capture regions (Supplementary Figure 4).

### Evaluation of PacBio HiFi Sequencing

We sequenced HG003, HG004, HG006, and HG007 with PacBio HiFi reads in order to contribute to Genome in a Bottle’s efforts to expand the confident regions. The high quality and mappability of PacBio HiFi data has recently proven instrumental in the first Telomere-to-Telomere human assembly[34] and undiagnosed rare disease[35]. Because we have sequence data for the same samples from PacBio HiFi and NovaSeq, we can quantify the differences in mappability and variant calling accuracy between the technologies.

We received ~21-fold PacBio HiFi coverage for HG003, HG004, and HG006, and ~29-fold for HG007, which were mapped with minimap2[36] and variants were called with DeepVariant v1.0 PacBio model. We compared callability at the exon and gene level, by setting a callability threshold of coverage between 5-fold and 150-fold. We used mosdepth[26] to quantify the fraction of bases in each RefSeq exon which met that criteria, and those which met the criteria when considering only reads with a mapping quality of more than 10 (Supplementary Figure5). We quantified RefSeq genes where any exon had less than 99% callability by either criteria (Figure 6 top left).

**Figure 6.**
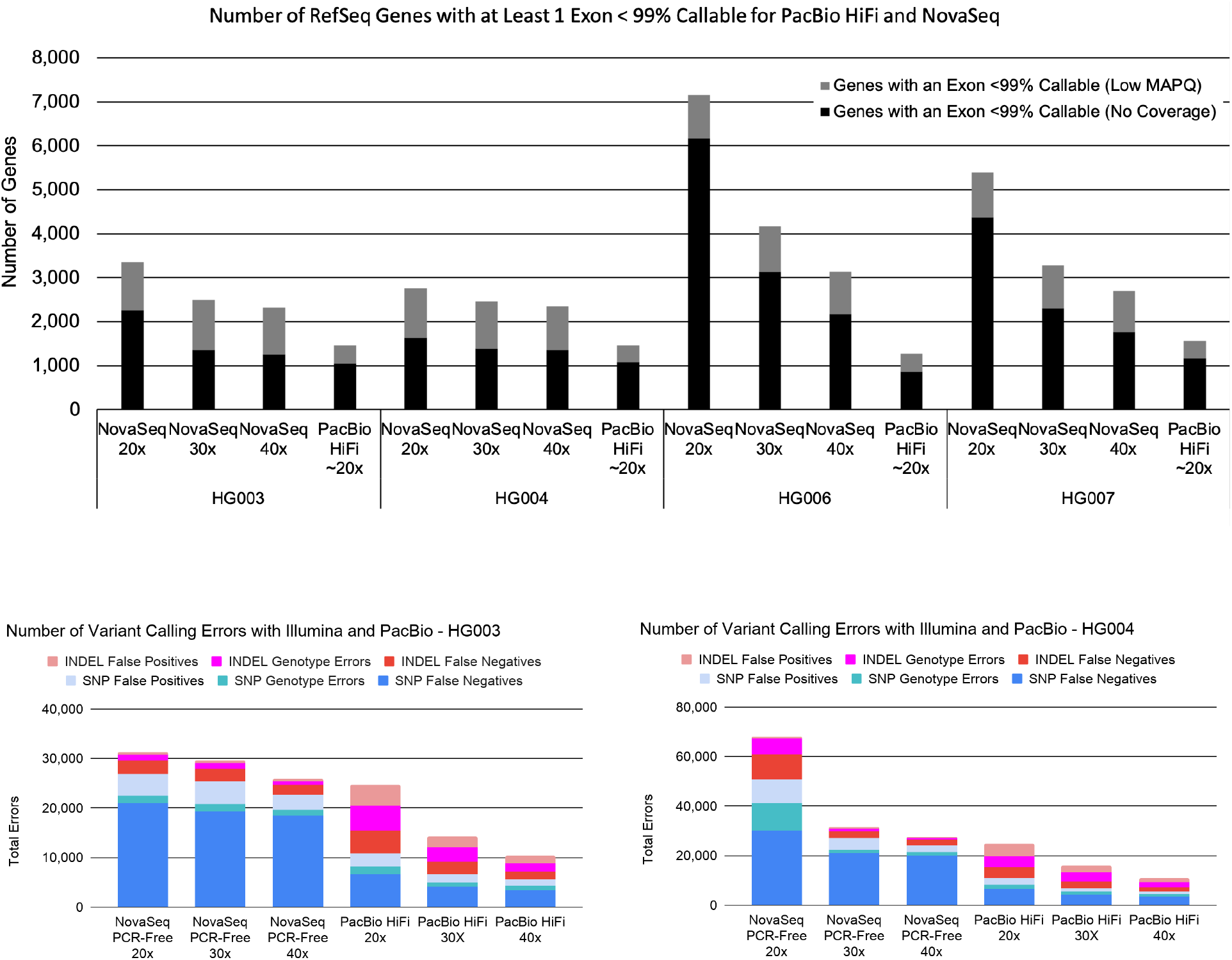
Comparison of PacBio HiFi and Illumina NovaSeq Data. Comparisons of the callability and accuracy of PacBio HiFi and PCR-Free NovaSeq WGS. The number of RefSeq genes which have any exon at less than 99% callability. This is defined by the number of bases covered to at least 5-fold coverage and less than 150-fold coverage by any read (black), or reads with a mapping quality more than 10 (grey) (top). The accuracy of mapping and calling variants with DeepVariant v1.0 using the Illumina model on NovaSeq samples or the PacBio model on PacBio HiFi samples for HG003 (bottom left) and HG004 (bottom right), as evaluated on the v4.2 Genome in a Bottle Truth Set. Note the difference in scale of the y-axis due to the larger number of errors in the HG004 NovaSeq 20x run.

PacBio HiFi sequencing at ~20x sequencing had fewer non-callable exons and genes than NovaSeq at 20x, 30x, or 40x coverage, when considering either bases at any coverage, or higher mapping quality bases. In addition, the PacBio HiFi callability metric was quite consistent across the four samples. We noted that the HG006 and HG007 NovaSeq samples had a much larger portion of genes with a non-callable base at 20x, and that increasing coverage in these samples reduced the variation between them with the other NovaSeq samples (HG003, HG004).

We assessed variant calling accuracy using the v4.2 Genome in a Bottle truth set for HG003 and HG004 samples. To understand the relationship between coverage and accuracy, for 30x and 40x coverage points we merged in additional PacBio HiFi data produced by Hudson Alpha for Genome in a Bottle. We observed that PacBio HiFi has fewer total variant calling errors for HG003 (Figure6 bottom left) and HG004 (Figure6 bottom right), driven by a large reduction in SNP false negative errors. This likely reflects the superior mappability of the long reads. Lower coverage for PacBio HiFi had more Indel errors, split roughly equally across false positives, false negatives, and genotype errors (calling a heterozygous event as homozygous and vice-versa). Indel errors are likely driven by difficulty determining homopolymer lengths.

The even split between these errors suggests that determining the correct call requires modeling the distribution in homopolymer lengths, with the optimal point balancing between undercalling and overcalling events. Although additional PacBio HiFi coverage had a modest positive effect on SNP accuracy, additional coverage substantially reduced Indel errors. Additional coverage of NovaSeq data had small reductions in False Negatives, while PacBio HiFi data continued to reduce errors at higher coverage points, suggesting that PacBio HiFi has a substantially higher ceiling of possible accuracy as coverage increases.

At the lower coverage points for NovaSeq, the HG004 sample had substantially more errors than the HG003 sample. This difference was reduced at higher coverages. A similar pattern - that of Illumina samples having higher variability at low coverages was seen with the HG006/HG007 mappability and with the analysis by component as well.

## Discussion

Our principal aim in this work is to provide the community with a dataset which is large enough to support method development efforts and training machine learning models, and representative enough to capture the various ways people sequence samples and standard run-to-run variability. This dataset is larger and more diverse than the one used in the first open-source release of DeepVariant, so we expect it can support the training of machine learning models of similar or greater complexity for production quality and use.

In the process of analyzing these data, we can distill a number of observations which may prove useful to those analyzing NGS data. We use this section to discuss these.

In our general coverage profiles, we noted certain samples with subtle differences (Figure2 top left). When stratifying with GA4GH regions, the differences were striking in certain regions (Figure2 top right). This suggests that the use of these GA4GH regions are useful to identify QC issues would otherwise not be as apparent in bulk analysis. We only observed these coverage issues in NovaSeq data, though there are not enough different runs to conclusively say this issue is more pronounced with the NovaSeq as opposed to a general issue.

In some cases, Illumina samples at lower coverage had more similar variant calling accuracy to the same sample at higher coverages (Figures 4,6). However, in other samples, there were large differences between low and high coverage points, with the low coverage point having much lower accuracy. Adding coverage seems to reduce inconsistency between samples, but is not always necessary for high accuracy. For labs wishing to balance cost and accuracy, this suggests a strategy where sequencing is initially performed at lower coverage and a number of QC methods are used to flag certain samples for additional sequencing. The GA4GH stratifications suggested above could be one such method.

We consistently observed a larger effect of coverage on Indel calling, compared to SNP calling. One possible explanation for this is that in larger tandem repeats or homopolymers, not all reads will span an event. Reads which do not span a repeat cannot be informative for insertions or deletions, and reduced coverage could make certain repeats uncallable.

Exomes had substantially greater variability in capture efficiency, mapped coverage, and downstream analysis accuracy. This suggests that reducing variability between runs is disproportionately beneficial for exome analysis, and that QC of exomes is more important. It may also suggest that informatics methods (calling, filtering, and post-processing) may benefit from tuning to the characteristics of the exomes in question. This highlights that though we speak of “exome sequencing” as a single term, the properties of two exomes can be very different. Analyses which use a cohort of multiple exomes (especially different capture types) will need to anticipate the batch effects that may arise.

The exome kit which performed best in this study is a property of the characteristics desired by the user. The Agilent v7 kit had the broadest capture across RefSeq, while the IDT xGen kit had the highest accuracy on the shared capture regions of all kits.

The PacBio HiFi samples obtained had a high level of consistency between runs, covered more exons and genes at greater than 99% callability, and even at 20x coverage had fewer errors, especially SNP false negatives. For samples where high accuracy across more of the genome is required, PacBio HiFi is uniquely capable.

It is important to note that the analyses presented here (mapping, coverage, and small variant calling) are a subset of the possible analyses one could conduct with exome and WGS sequencing. It is possible that looking at structural variant calling, copy number analysis, RNA expression, somatic calling, or other methods could reveal a large difference between instruments or sample preparation methods that are not visible here. We hope that the sequence data released in this study will prove useful to experts in fields beyond the ones investigated in detail here.

## Methods

### Generation of Sequencing Data

We contracted HiSeqX, NovaSeq, and PacBio HiFi sequencing from Novogene (https://en.novogene.com/). All WGS and exome runs were conducted with 151-bp paired-end reads. The instructions provided with the order was for 50x coverage WGS (which involved a higher cost than for standard 30x) of PCR-Free and PCR-Positive samples for HG001-HG007, NA12891, and NA12892.

For exomes, we asked Novogene for the three most commonly used capture kits, which were indicated to be Agilent v7, IDT-xGen, and Nextera. We requested 200x coverage for each capture kit from the NovaSeq and HiSeq4000 platforms. To quantify deduplicated, mapped coverage in exomes, we performed the calculator only only the capture regions of the exome kit. Because we could only find capture regions for these exome kits on GRCh37, determination of exome coverage was determined on the GRCh37 reference, specifically hs37d5.

For PacBio HiFi data, we requested 3 SMRT Cells 8M for each sample of HG003, HG004, HG006, and HG007. Libraries were prepared targeting a 15kb insert size, and sequenced on Sequel II System with Chemistry 2.0.

For physical samples for NA12891, NA12892, HG006, and HG007, we contracted Novogene to procure samples from Corriell. Samples for HG001-HG005 were generously provided by NIST Genome in a Bottle from their reference standard collection.

### Downsample and Mapping

After receipt, samples were mapped to GRCh38 with BWA MEM [21] in an ALT-aware manner and deduplicated with Picard MarkDuplicates [24]. Variants were called by DeepVariant v1.0 [12] and the median coverage at candidate positions taken as the mapped, deduplicated coverage for the files. For exomes, the same process was performed only on the capture regions for each kit.

Downsample fractions to reach 40x, 30x, and 20x coverages for WGS from these values were calculated from each sample. FASTQ files were then downsampled using SEQTK [37]. For exomes, the same process was performed with coverage targets of 100x, 75x, and 50x.

### Coverage and Mapping Statistics

Statistics for rate of duplicate, unmapped, supplementary, and off-target reads were calculated with samtools flagstat using samtools v1.9 [25]. Base-level coverage statistics for whole genome, exome capture regions, and for exon/gene level statistics were generated with mosdepth v0.2.9 [26].

### Variant Calling

Variant calling was performed using DeepVariant v1.0, Strelka v2.9.10, Octopus v0.6.3-beta, and GATK versions 4.1.8.0 and 4.1.8.1. The commands used for each caller are included in the supplementary material. Though different GATK versions were used for different data types, we do not expect this to have a major effect on the results.

Call sets were assessed using v0.3.9 of the haplotype comparison tool, hap.py. The v4.2 truth sets from GIAB were used to benchmark HG002-4 samples mapped to GRCh37, and the v.3.3.1 truth sets were used for HG001-7 samples mapped to GRCh38. For exomes, evaluation was performed only on regions representing the intersection of all capture kits.

### PacBio HiFi Sequencing

In addition to releasing this data here, we previously released this data to Genome in a Bottle (ftp://ftp-trace.ncbi.nlm.nih.gov/giab/ftp/data/AshkenazimTrio/HG003_NA24149_father/PacBio_CCS_Google_15kb/) and to the Human Pangenome Reference Consortium, where it is indexed (https://github.com/human-pangenomics/HG002_Data_Freeze_v1.0) with a mirror on AWS S3.

## Supporting information

Supplementary File 1. Download links for FASTQ files

Supplementary File 2. Download links for BAM files

Supplementary File 3. Download links for VCF files

## Acknowledgements

This work depends on the exceptional effort of the National Institutes of Standards and Technology in establishing the Genome in a Bottle truth sets. Samples for HG001-HG005 were generously provided by the National Institutes of Standards and Technology to Novogene for sequencing. We also thank the participants in the Personalized Genomes Project who contributed their DNA and cell lines for the truth set samples. This contribution was essential to this work, as well as to broad advances in the development of reliable genome analysis methods. This work relies on the open-source contributions of a variety of methods in read mapping, QC, variant calling, and accuracy assessment. We thank the authors and maintainers of all of these tools.

We thank Justin Zook and Nate Olson from Genome in a Bottle, Aaron Wenger and Billy Rowell from Pacific Biosciences, and Nick Furlotte from Google for comments and suggestions on the manuscript.

Sequencing instruments were run at Novogene. We thank the scientists who conducted these sequencing runs for their efforts which generated the data in this work.

## Author contributions

AC, PC, and HY conceived the study. AC and PC designed the study. AC and HY procured sequence data. GB, PC, MN, AK, SG built infrastructure supporting rapid and robust analysis. AC and GB performed experiments and analyzed results. AC and GB wrote the manuscript.

## Data availability

All data are released under a CC-0 license. The sequence files are publicly available in Google Cloud Storage in the gs://brain-genomics-public/research/sequencing/

(console.cloud.google.com/storage/browser/brain-genomics-public/research/sequencing/). Instructions for accessing this public data can be found in Google Cloud Storage documentation (cloud.google.com/storage/docs/access-public-data).

---

**Supplementary Figure 1.**
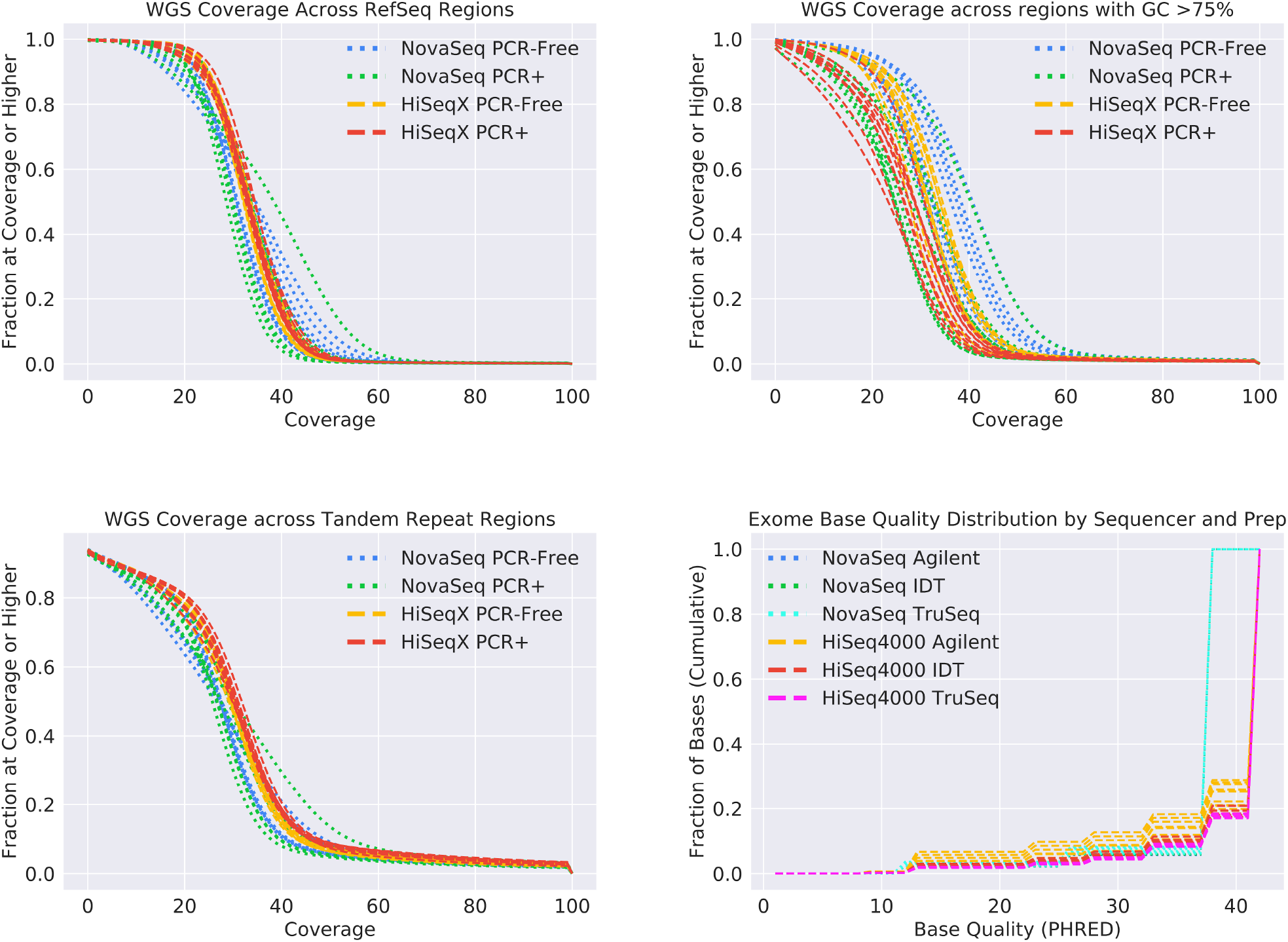
Additional genome coverage and read quality attributes for samples in this study. The fraction of the genome in each WGS sequencing run covered to a least a given coverage, analyzed for various genome regions. Subsetting analysis to regions annotated as exon by RefSeq (top left), for regions annotated as having GC content >75% in the GA4GH region stratifications (top right), for regions annotated as tandem repeat or homopolymer in the GA4GH region stratifications (bottom left). The base quality distribution by sequencer and preparation for the exome runs for all reads, including on-target, off-target, and unmapped (bottom right).

**Supplementary Figure 2.**
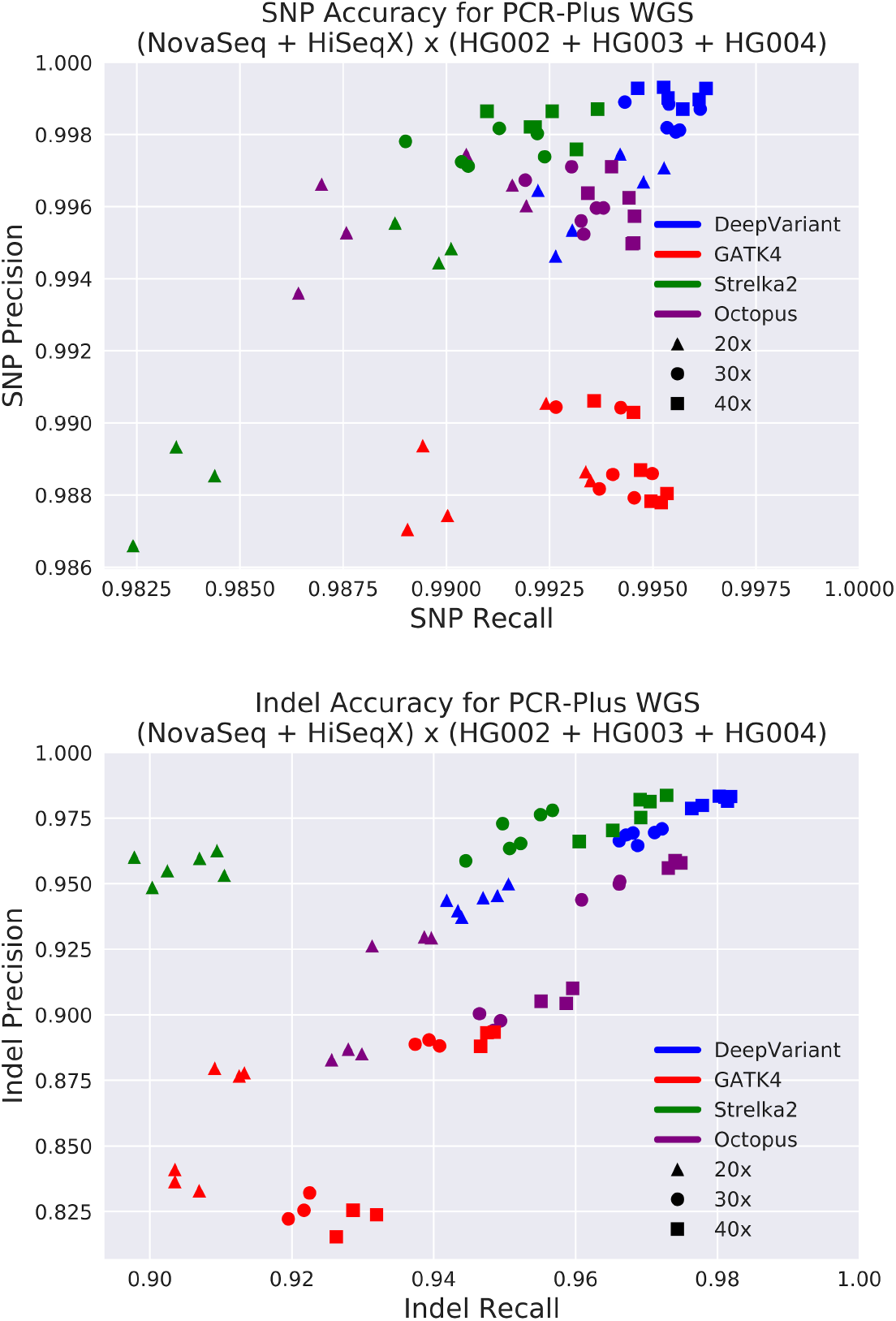
Variant Calling Accuracy for PCR-Positive WGS Samples. Precision and Recall for SNP (top) and Indel (bottom) for PCR-Positive WGS sequence runs, compared with the v4.2 GIAB truth set on BAM files mapped to GRCh38. All accuracies are for chromosome20, which is not used for training DeepVariant. Accuracies across multiple PCR-Positive runs of HiSeqX and NovaSeq for each of HG002, HG003, and HG004 at 20x, 30x, and 40x coverage (6 points at each coverage, 18 total runs). Four different callers were run on the same samples, coded by color: DeepVariant, GATK4, Strelka2, Octopus.

**Supplementary Figure 3.**
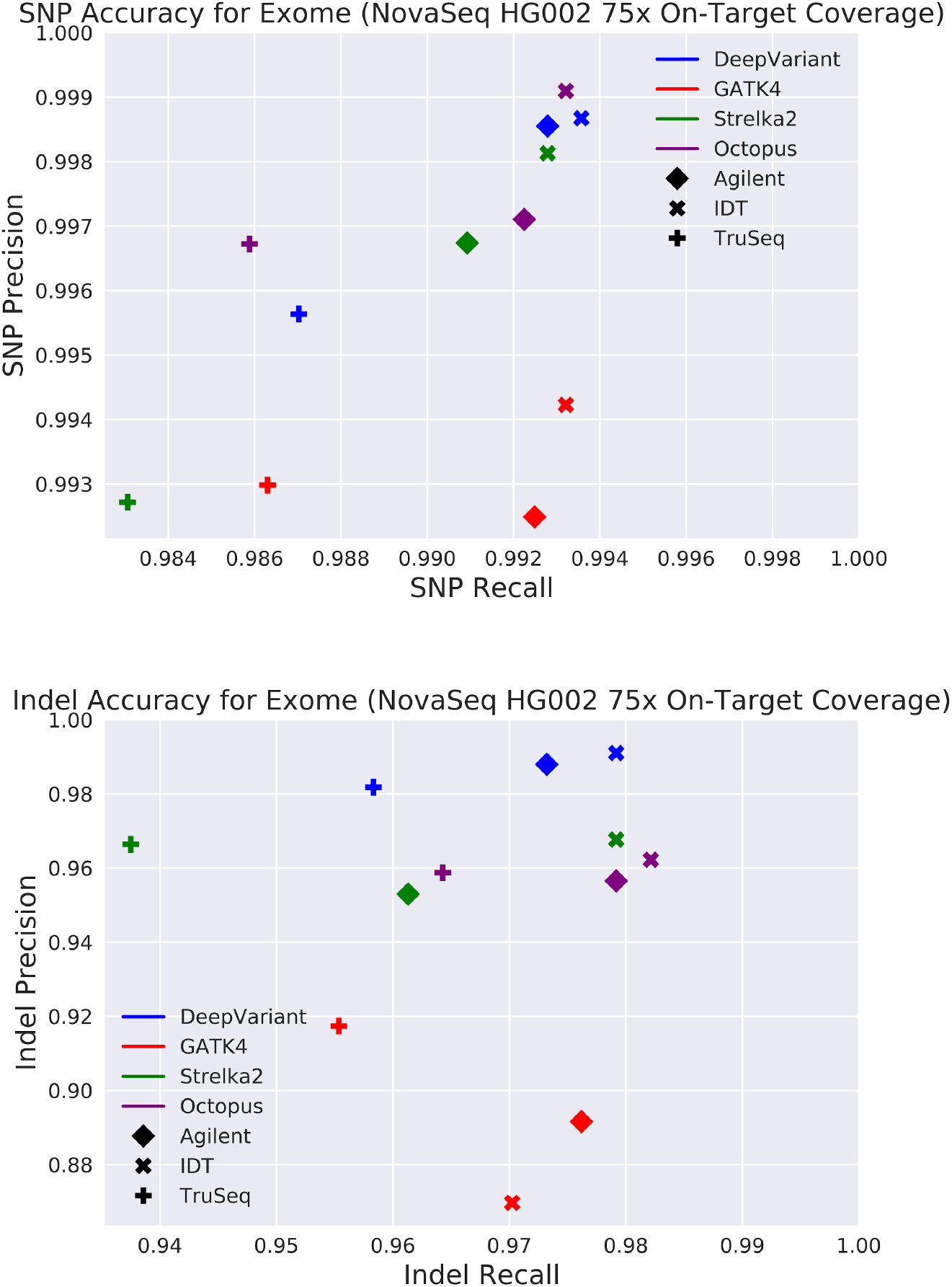
Variant Calling Accuracy for Exome Kits at a Single Instrument, Sample, and Coverage. Precision and Recall for SNP (left) and Indel (right) for exome sequence runs, compared with the v4.1 GIAB truth set on BAM files mapped to GRCh37 and evaluated on the common intersection of capture regions. All accuracies are across the full genome in order to have enough variants for a stable comparison. Four different callers were run on the same samples, coded by color: DeepVariant, GATK4, Strelka2, Octopus, with capture kits coded by shape for: Agilent, IDT, TruSeq. In this case a single sample (HG002), instrument (NovaSeq), and coverage (the middle 75x point) are shown in isolation to illustrate how differences between kit and caller look on an exemplar. This is similar to the bottom four panels of Figure 3 for WGS, but for exomes.

**Supplementary Figure 4.**
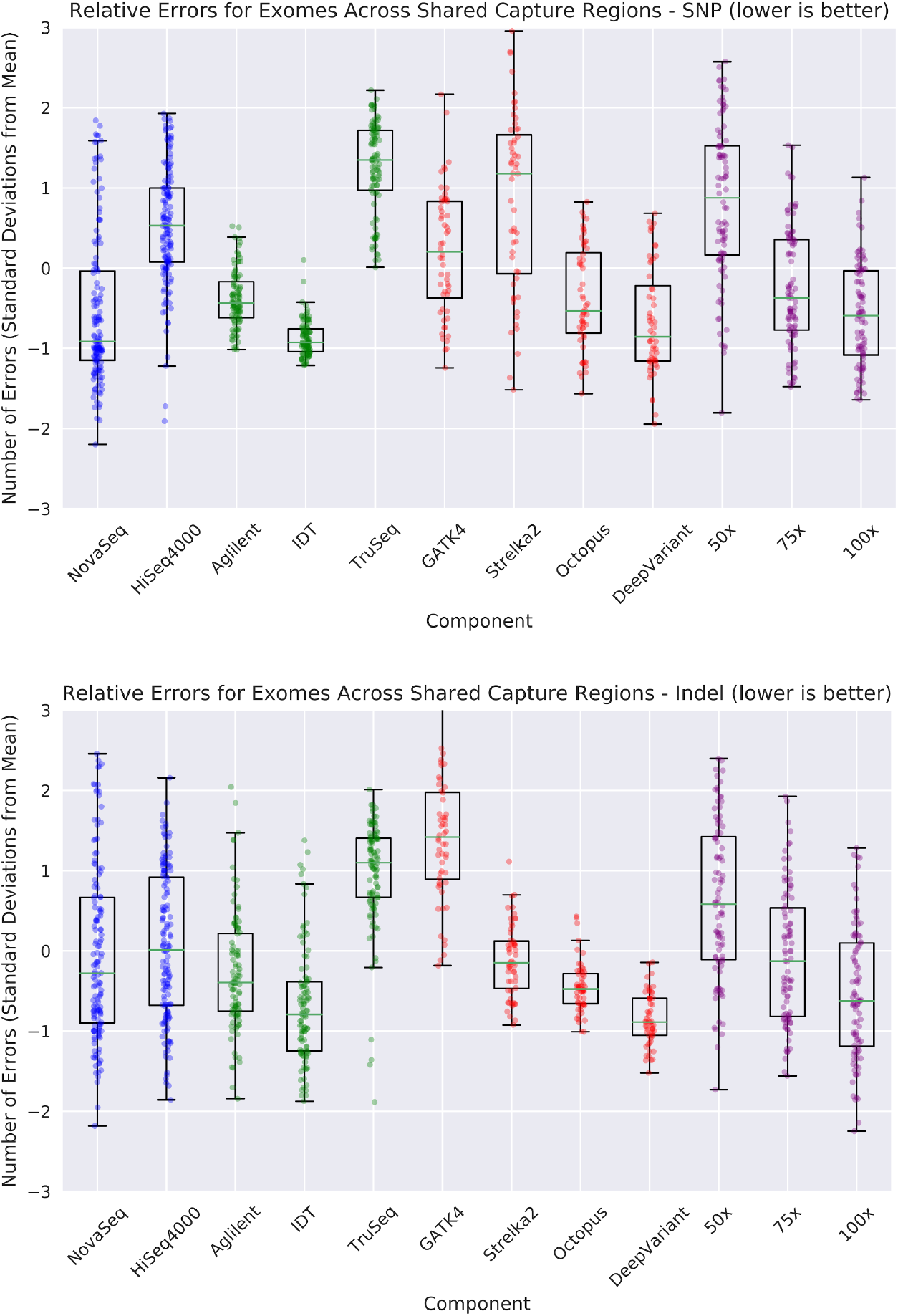
Contributions of Components to Exome On-Target Variant Calling Pipeline Accuracy. Measuring the contributions of component: instrument (red), preparation/kit (green), variant caller (red), and coverage (purple) to SNP (left) and Indel (right) errors in variant calling. The mean and standard deviation for error is calculated, and for each sample the number of standard deviations from the mean of errors is plotted. For example, every dot in the NovaSeq column is a run using NovaSeq data where there is another run with all of the same properties except the use of HiSeq4000 in the HiSeq400 column. When a dot is below y-axis=0, it means the use of that factor reduces error relative to the “average” run. This is shown for exome pipelines considering only the shared intersection of capture regions for SNPs (top) and Indels (bottom).

**Supplementary Figure 5.**
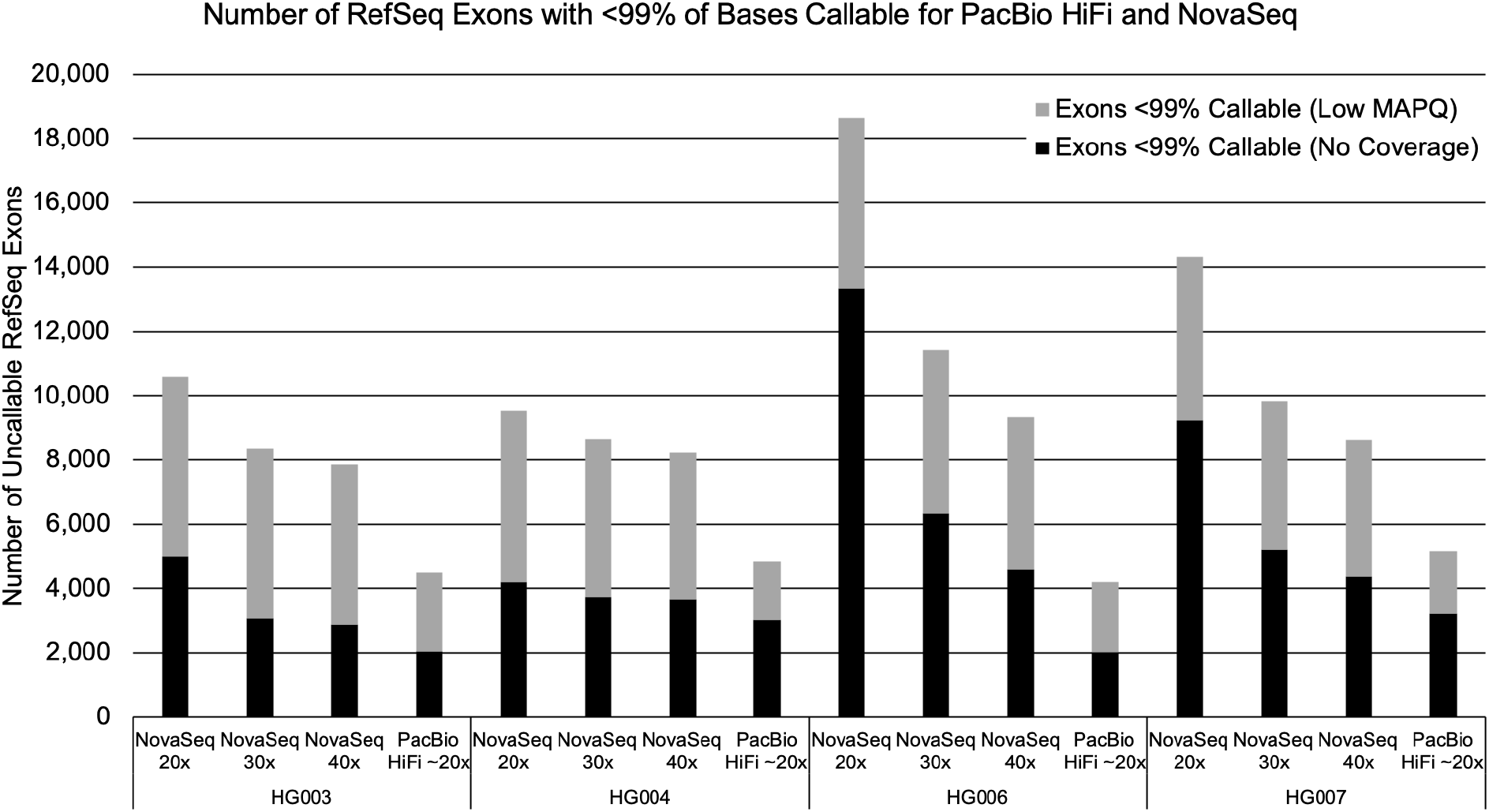
Comparison of Exon-Level Callability Between PacBio HiFi and Illumina NovaSeq. Comparisons of the callability and accuracy of PacBio HiFi and PCR-Free NovaSeq WGS. This shows the number of RefSeq exons with less than 99% callability. This is defined by the number of bases covered to at least 5-fold coverage and less than 150-fold coverage by any read (black), or reads with a mapping quality more than 10 (grey).

## Reference Genomes Used

GRCh37 - hs37d5

ftp://ftp-trace.ncbi.nih.gov/1000genomes/ftp/technical/reference/phase2_reference_assembly_sequence/hs37d5.fa.gz

GRCh38 - GRCh38.p13

https://www.ncbi.nlm.nih.gov/assembly/GCF_000001405.39/

GRC38_no_alt

ftp://ftp.ncbi.nlm.nih.gov/genomes/all/GCA/000/001/405/GCA_000001405.15_GRCh38/seqs_for_alignment_pipelines.ucsc_ids/GCA_000001405.15_GRCh38_no_alt_analysis_set.fna.gz

## Commands Used

### BWA

~~~
bwa mem -t ${THREADS} “${REF}” “${READS}” “${MATES}” -R
“@RG\\tID:${RGID}\\tPL:ILLUMINA\\tPU:NONE\\tLB:${RGID}\\tSM:${SAMPLE}” |
samtools sort -O BAM -o ${BWA_OUTPUT}
~~~

### MarkDuplicates

~~~
java -jar gatk-package-4.1.2.0-local.jar MarkDuplicates -I ${BWA_OUTPUT} -O
${OUTPUT} -M ${OUTPUT%.bam}.metrics
~~~

### Mapping and Coverage Statistics

~~~
samtools flagstat ${BAM} > ${BAM%.bam}.flagstat.txt
mosdepth -n -x --quantize 0:15:150: mosdepth.${BAM%.bam} ${BAM}
mosdepth --by grch38_refseq.bed mosdepth.${BAM%.bam}.capture ${BAM}
mosdepth -n -b hg38.refseq.gene_names.bed -T 0,5,150
mosdepth.${BAM%.bam}.refseq ${BAM}
~~~

### DeepVariant

WGS:

~~~
sudo docker run \
  -v “${INPUT_DIR}”:”/input” \
  -v “${OUTPUT_DIR}:/output” \
  google/deepvariant:”${DV_VERSION}” \
  /opt/deepvariant/bin/run_deepvariant \
  --model_type=“${MODEL_TYPE}” \
  --ref=/input/”${REF}” \
  --reads=/input/”${READS}” \
  --output_vcf=/output/output.vcf.gz \
  --num_shards=“$(nproc)” \
  --make_examples_extra_args=“downsample_fraction=${DOWNSAMPLE_FRACTION}” \
  --output_gvcf=/output/output.g.vcf.gz
~~~

For WES, the following flag was added:

~~~
  --regions=/input/”${CAPTURE_BED}”
~~~

### GATK HaplotypeCaller

WGS:

~~~
sudo docker run \
  -v “${INPUT_DIR}”:”/input” \
  broadinstitute/gatk gatk CreateSequenceDictionary \
  -R /input/”${REF}” \
  -O /input/”$(basename “${REF}” .fa).dict”
sudo docker run \
  -v “${INPUT_DIR}”:”/input” \
  -v “${OUTPUT_DIR}”:”/output” \
  broadinstitute/gatk gatk HaplotypeCaller \
  -R /input/”${REF}” \
  -I /input/”${READS}” \
  -O /output/output.vcf.gz \
  --native-pair-hmm-threads “$(nproc)”
~~~

For WES, the following flag was added to the second command:

~~~
   -L /input/”${CAPTURE_BED}”
~~~

### Octopus

WGS:

~~~
venv/bin/octopus \
  -R “${REF}” \
  -I “${READS}” \
  -o “${OUTPUT_DIR}/output.vcf.gz” \
  --threads
~~~

For WES, the following flag was added:

~~~
   -t “${CAPTURE_BED}”
~~~

### Strelka2

WGS:

~~~
sudo “${STRELKA_INSTALL_PATH}”/bin/configureStrelkaGermlineWorkflow.py \
  --bam “${READS}” \
  --referenceFasta “${REF}” \
  --runDir “${OUTPUT_DIR}”
sudo “${OUTPUT_DIR}”/runWorkflow.py \
  -m local \
  -j “$(nproc)”
~~~

For WES, the following flags were added to the first command:

~~~
  --callRegions “${CAPTURE_BED}” \
  --exome
~~~

### Haplotype Comparison with hap.py

Base command for WGS and WES:

~~~
sudo docker run -i \
  -v “${INPUT_DIR}”:”/input” \
  -v “${OUTPUT_DIR}”:”/output” \
  pkrusche/hap.py /opt/hap.py/bin/hap.py \
  /input/”${TRUTH_VCF}” \
  /output/output.vcf.gz \
  -f /input/”${TRUTH_BED}” \
  -r /input/”${REF}” \
  -o /output/happy.output \
  --engine=vcfeval
~~~

For all WGS and some WES runs, evaluation was performed only on chromosome 20 by adding the following flags. EVAL_REGION was set to either chr20 or 20, depending on the reference genome used.

~~~
  -l “${EVAL_REGION}”
~~~

For WES, the following flag was added:

~~~
  -T /input/”${CAPTURE_BED}”
~~~

